# The application of neural networks to classify dolphin echolocation clicks

**DOI:** 10.1101/2022.06.14.496047

**Authors:** Vahid Seydi, Lucille Chapuis, Gemma Veneruso, Sudha Balaguru, Noel Bristow, David Mills, Lewis Le Vay

**Affiliations:** Centre for Applied Marine Sciences, School of Ocean Sciences, Bangor University, Menai Bridge, United Kingdom

## Abstract

Passive acoustic monitoring (PAM) is a common approach to monitor marine mammal populations, for species of dolphins, porpoises and whales that use sound for navigation, feeding and communication. PAM produces large datasets which benefit from the application of machine learning algorithms to automatically detect and classify the vocalisations of these animals. We present a deep learning approach for the classification of dolphins’ echolocation clicks into two species groups in an environment with high background noise. We compare the use of Convolutional Neural Networks (CNN) and Recurrent Neural Network (RNN), in which we feed the models the raw waveform data and spectrograms. We show that both models perform well, with the highest performance achieved by a CNN fed with spectrograms (F1 score 97 %) and an RNN fed with raw data (F1 score 96%) fitted with Gated Recurrent Units (GRU). We recommend the use of such models to classify echolocation clicks in marine environments where background noise levels exhibit high spatial and temporal variance. In particular, the RNN showed excellent performance, while being fed with raw data, in terms of reduced processing time and storage. Deep learning automatically extracts effective features from the raw waveform in the training process through multiple layers of the model, without the need to rely on feature extraction in a separate pre-processing step.

## 1. INTRODUCTION

Passive acoustic monitoring (PAM) is a popular method used for monitoring odontocetes (toothed whales) in their ecosystems (Mellinger et al., 2007). Odontocetes produce series of clicks, whistles, and other complex sounds to survey their surroundings, hunt for food, and communicate with each other. Recording these signals with hydrophones allows the examination of species occurrence, distribution, density, behaviour, and the consequences of disturbance at specific spatial and temporal scales (Booth et al., 2020; Mellinger et al., 2007). Such information is critical to monitor and understand populations of odontocetes, which are otherwise difficult to monitor, and for which some species are declining and/or under threatened conservation status (Nelms et al., 2021).

In particular, delphinids (dolphins) produce clicks with their biosonar system and utilise their returning echoes to find and identify prey and discern environmental structure. Clicks are impulsive short pulses of substantial strength, lasting for microseconds. Delphinids produce multiple clicks in a row with diverse cue rates ranging from 0.5 to 2 clicks per second (Au, 1993). Clicks from delphinids are important vocalisations, predominant in comparison with whistles or other calls, and are used for navigation and prey detection (Au et al., 2000). Therefore, detecting and identifying delphinid clicks is an important aspect of PAM applications, for example when monitoring cetacean interactions with offshore infrastructure developments. In particular, the development and future operation of marine renewable energy (MRE) infrastructure raises the question of potential interactions of local populations of delphinids with individual turbines and turbine arrays, making the determination of the presence or absence of key species of critical importance if harm to cetaceans is to be avoided (see e.g., Gillespie et al., 2021; Palmer et al., 2021). Similarly, the development of delphinid deterrents to avoid by-catch or interactions with anthropogenic structures requires accurate monitoring procedures to assess their specific performance (Schakner and Blumstein, 2013). Spectral and temporal properties of delphinid echolocation clicks have been investigated to help classify these vocalisations to species (Roch et al., 2011; Soldevilla et al., 2008). Clicks are typically classified based on a finite number of features, such as their spectral content, duration and inter-click intervals. However, interspecific patterns are not always identified, and the intraspecific variability of the clicks make any generalisation difficult. More recently, automatic methods have been proposed to automatically classify clicks produced by whales and dolphins, with mixed results (Bermant et al., 2019; Frasier et al., 2017; Gillespie and Chappell, 2002; Griffiths et al., 2020; Jarvis et al., 2008; Luo et al., 2017; Roch et al., 2008, 2011).

With the advancement of data storage capabilities and requirement for long-term monitoring, PAM can generate terabytes of data, and manual classification of delphinid clicks is time-consuming and may be inconsistent among analysts. For the last decade, machine learning techniques have provided improved detection and classification methods for PAM of cetaceans, with varying performance outputs (Usman et al., 2020). The cetacean vocalisations are usually pre-processed via the extraction of features, followed by the detection and classification stages. Furthermore, these stages are commonly processed retrospective to the collection of the recordings, as they do not allow real-time processing.

Deep neural networks are the extension of shallow artificial neural networks that allow a machine to be fed with raw data and to automatically discover the representations, or features, needed for detection or classification (LeCun et al., 2015). Convolutional neural networks (CNN) and recurrent neural networks (RNN) are two popular structures in Deep learning models. Although primarily used in visual recognition contexts, CNNs are becoming more and more widespread for audio-related tasks (Piczak, 2015; Valenti et al., 2017), and have been used successfully to detect and classify marine mammal vocalisations (Allen et al., 2021; Bermant et al., 2019; Ibrahim et al., 2021; Shiu et al., 2020; Zhong et al., 2020). In parallel, RNNs are excellent for the extraction of patterns in time series or signals (LeCun et al., 2015) and have recently been used to classify sperm whale clicks to recognise vocal clans and individual whales (Bermant et al., 2019). Delphinid clicks have been previously classified successfully by unsupervised neural networks (Frasier, 2021; Frasier et al., 2016, 2017), suggesting the ability of these networks to recognise different patterns and/or features in echolocation clicks. However, the classes could not all be identified as a taxonomic group (ie. species), and the complex workflow included many succeeding steps of clustering, labelling and classification (Frasier, 2021).

Generally, detection and classification tasks are affected by the amount of background noise present at the recording sites, and the varying signal-to-noise ratio when considering marine mammal vocalisations. For example, tidal sites are affected by periods of high current speeds, which significantly increase the background noise, sometimes with a difference of more than 30 dB re 1 μPa between the tidal phases (Willis et al., 2013). Similarly, underwater structures and machinery, like tidal turbines, offshore windfarms or other MRE installations, and other anthropogenic activities can also alter background noise in regular or irregular patterns. PAM detection and classification of animals’ signals in such environments may suffer from a low signal-to-noise ratio and in the case of machine learning models, a low generalisation performance.

In this study, we investigate the use of CNN and RNN to efficiently extract the informative features from delphinid clicks, recorded at a strong tidal flow site with an MRE infrastructure in place, for classification and similarity analysis. We test both methods by feeding the raw data (click waveforms) and spectrograms (image representation of the clicks) to the networks. Clicks from three species of dolphins common to the region were collected, labelled and used to train four different models: (i) CNN with raw data; (ii) CNN with spectrograms; (iii) RNN with raw data; (iv) RNN with spectrograms. The models were used to automatically classify the clicks into two groups: Risso’s dolphin (*Grampus griseus*) clicks, and delphinid clicks from species that produce broadband clicks, combining clicks from common (*Delphinus delphis*) and bottlenose dolphins (*Tursiops truncatus*). The combination of click data for these two species (bottlenose and common dolphins) is typical for traditional PAM studies, as they are not easily distinguished by human operators due to the similarities in click properties (Au, 2002), hence the training of supervised machine learning models is not currently possible.

## 2. METHODS

### 2.1. Description of PAM system and deployment

The PAM system was deployed in Wales, UK, in Holyhead Deep (53° 17.796 ‘ N, 4° 47.948’ W), at 85 m depth for one month in total (August – September 2019). The site included an MRE infrastructure which typically produced anthropogenic noise, notably due to the movement of metallic components, such as mooring gear including chains. A Sonar Point recorder (Desert Star Systems, USA) was connected to four HTI 99-UHF hydrophones (HTI, High Tech Inc, USA), recording at 312,500 samples per second. Only one data channel (i.e. one hydrophone) was used for analyses in this study. The change in background noise was investigated by calculating the RMS (root-mean-squared) sound level for each hour of recordings and is presented in supplementary material Fig. S1.

### 2.2. Click detection and data labelling

PAMGuard (Gillespie et al., 2009; www.pamguard.org) was used to process the acoustic data and detect all clicks. The click detector was configured to be triggered by transient signals with peak frequencies above 20 kHz that rose 10 dB above a continuous measure of background noise. When triggered, the detector stores short clips (~ 1 ms) of unfiltered data. An experienced analyst (GV) then manually classified 16,520 clicks into Risso’s dolphin clicks (10,367) and broadband click species events (6,153), based on the click frequency characteristics (peak frequency and bandwidth) and waveforms (Palmer et al., 2017; Soldevilla et al., 2008).

### 2.3. Feature extraction

Two approaches were considered to prepare the data for the deep learning models. In the first approach, we used the ability of neural networks to learn inherent rules from the input raw dataset: the model automatically extracts the appropriate features required to classify the data. In the second approach, the click frequencies are extracted using a short-time Fourier transform (STFT) and given to the model in the form of a spectrogram. In this approach, the STFT window size and the distance between adjacent windows are considered as hyperparameters. In the hyperparameter tuning process, varying values were examined to find near-optimal values for both.

### 2.4 Machine learning classifier

We tested two popular classification models in the deep learning domain, CNN and RNN, to classify acoustic signals (Usman et al., 2020). The main characteristic of CNN models is automatic feature extraction by applying convolution filters based on local correlations. On the other hand, RNN models are specialised for processing a sequence of values and use outputs from past or future inputs of the temporal sequence to inform the current prediction (Goodfellow et al., 2016). However, standard RNN cannot work efficiently with long-term dependencies in data where the distance between the relevant information and the place where it is required is large. Therefore, a Long Short-Term Memory (LSTM) network, a special case of RNN that takes long-term dependencies into account, was tested, as well as a Gated Recurrent Unit (GRU) network, which represents a moderate version of recurrent gate, and has been used previously for human speech recognition (Shewalkar et al., 2019).

As with most supervised learning tasks, the process comprises three phases: (1) the training phase in which the weights or parameters of the model are trained on labelled data, (2) the validation phase in which the hyper parameters are tuned, and (3) the test phase where the model is evaluated as a classifier for unseen labelled data. In the test and validation phases, the model parameters do not change, and the model is evaluated based on their achieved values. Therefore, these two phases are also referred to as the evaluation phase.

### 2.5 Convolution Neural Networks architecture

Two models of CNN were used, depending on the input type: we used one-dimensional convolution filters on the raw data and two-dimensional convolution filters on the spectrogram dataset. As shown in Figure 1, a CNN consists of the following blocks:

a. The **Input layer**, is the raw signal waveform for one-dimensional convolutional networks and is the spectrogram for two-dimensional convolutional networks. The dimensions of the spectrograms are determined by the size of the STFT window and the distance between adjacent windows.
b. The **Convolution layer** consists of several convolution filters (1D or 2D). They represent weights or model parameters, whose optimal values are attained in the training process. The dimension and stride of the filters are considered as hyperparameters of the model and their optimal value is obtained in the validation process. Convolution filters play the key role in the feature extraction process.
c. The **Batch Normalisation layer** keeps the mean output close to 0 and the output standard deviation close to 1.
d. The **Activation function** is a nonlinear function that takes the output of the batch normalisation layer as input, aiming to capture the nonlinearity of the samples. ReLU and Leaky-ReLU are two popular activation functions which are traditionally used in most CNN models.
e. The **Pooling layer** has the role of extracting the most important features from the extracted features of the convolution layer. In this layer, a down-sampling mechanism is performed by averaging or maximising. The size of filter and stride are considered as hyperparameters.
f. The **Fully Connected layers** apply a weighted linear mapping so that the features move closer to the target space.
g. The **SoftMax or Logistic layer** is the last layer of CNN and appends at the end of the fully connected layer. The Logistic class and the SoftMax class are chosen for binary classification and multi-class classification, respectively. Since our problem is binary classification, Logistic is used in the proposed CNN model.

**Figure 1:**
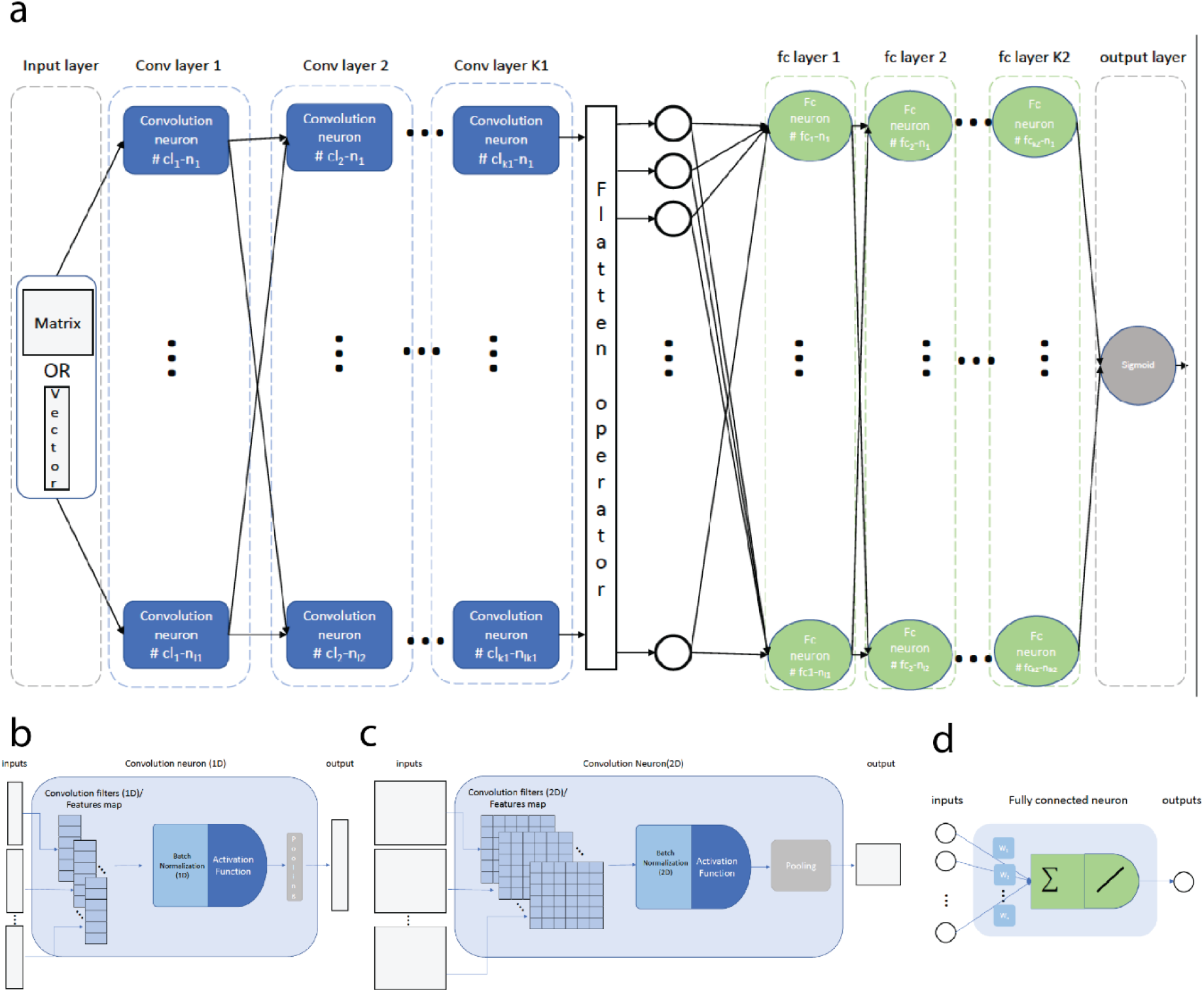
Architecture of the Convolutional Neural Network: a) general view of the CNN model b) enlarged view of a one-dimensional convolution neuron c) enlarged view of a twodimensional convolution neuron d) enlarged view of a fully connected neuron.

All the hyperparameters used in both CNN models are summarised in Table 1.

**Table 1:** Description of the four models tested and their hyperparameters.

| Model | Category | Parameters | Values |
| --- | --- | --- | --- |
| 1. CNN,<br>raw signals | Convolution Blocks | Number of CNN blocks | 4 |
|  |  | Number of channels | [16, 32, 64, 48] |
|  |  | Convolution filter size | [5, 5, 9, 5] |
|  |  | Convolution filter stride | [1, 3, 1, 1] |
|  |  | Activation function | ReLU |
|  |  | Max-pool filter size* | [2, 3, 1, 2] |
|  |  | Max-pool stride | 1 |
|  | Linear blocks | number of fully connected layers | 4 |
|  |  | Number of neurons | [512, 512, 256, 64] |
|  | Others | Drop out probability | 0.4 |
|  |  | Learning rate | 1e-4 |
| 2. CNN,<br>spectrograms | Spectrogram | n-fft | 128 |
|  |  | Hop size | 32 |
|  | Convolution Blocks | Number of blocks | 5 |
|  |  | Number of channels | [48, 128, 64, 16, 32] |
|  |  | Convolution filter size | [3, 3, 3, 3, 3] |
|  |  | Convolution filter stride | 1 |
|  |  | Activation function | ReLU |
|  |  | Max-pool filter size* | [1, 1, 1, 2, 1] |
|  |  | Max-pool stride | 1 |
|  | Linear blocks | number of layers | 4 |
|  |  | Number of neurons | [1024, 64, 64, 64] |
|  | Others | Drop out probability | 0.2 |
|  |  | Learning rate | 1e-4 |
| 3. RNN,<br>raw signals | Neural network | Number of layers | 3 |
|  |  | Number of neurons | 60 |
|  |  | Type of gate | GRU |
|  |  | Learning rate | 0.000723 |
| 4. RNN,<br>spectrograms | Spectrogram | n-FFT | 16 |
|  |  | Hope size | 8 |
|  | Neural network | Number of layers | 4 |
|  |  | Number of neurons | 40 |
|  |  | Type of gate | LSTM |
|  |  | Learning rate | 0.000975 |

\* for max-pool filter size, number 1 means not using max-pool filter in that block

### 2.5 Recurrent Neural Networks architecture

The general structure of an RNN can be seen in Figure 2. A sample is divided into a sequence of consecutive states. The states are given to the first layer of the model, which involves several RNN/GRU/LSTM gates (neurons), one after the other. The standard RNN gate is a basic neuron which is fed with two inputs: one directly from the input and the other one comes from the output of its previous state. Depending on how long history can be captured, two general extensions have been utilised to the structure of the RNN neuron: the GRU gate and the more complex LSTM gate. To add more flexibility to the model several recurrent layers can be stacked. A Fully Connected layer and a Logistic layer reside at the end of recurrent layers to transform extracted features to the target label. In this way, the history of the input sequence is considered to extract features.

**Figure 2:**
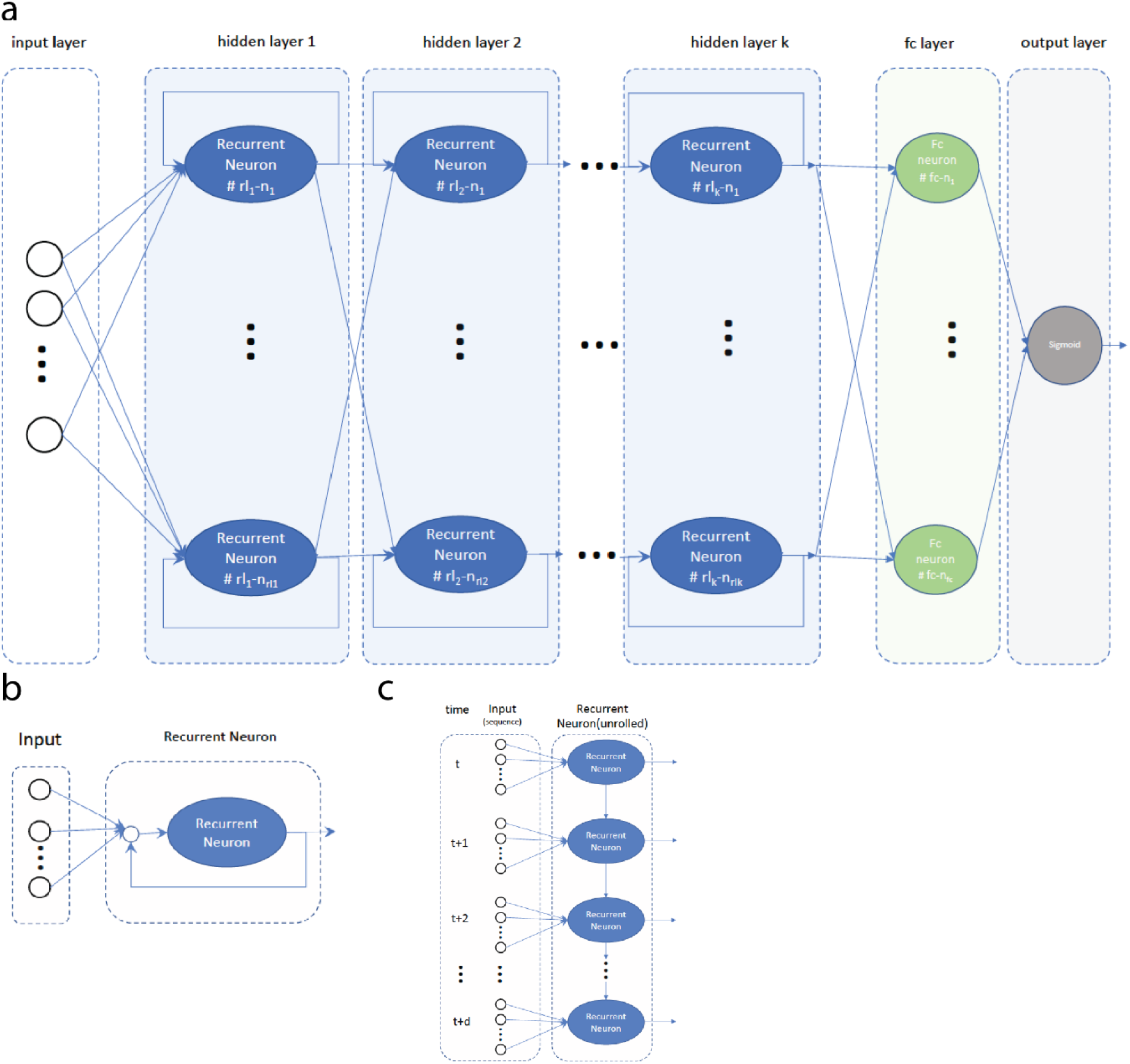
Architecture of the Recurrent Neural Network: a) General view of the RNN b) enlarged view of a recurrent neuron c) enlarged view of an unrolled recurrent neuron.

In this structure, the number of gates in each layer, the number of recurrent layers, and the type of gates are considered as hyperparameters. Also, since we used two different forms of input, we used two distinct methods to transform a sample into a sequence of states, as follows:

In the case that the inputs are given to the model in the form of a raw wave with N sample points, n (n ≤ N) consecutive samples are considered as a state, n referring to the dimension of input. Therefore, each sample contains N/n states (the length of sequence). Since the maximum click length is 512, the value of N is 512 and n was considered as a hyperparameter so that the near-optimal value could be achieved.
In the case of spectrograms as inputs, each column of the matrix, which is the magnitude of frequency coefficient of the click signal in a window of STFT, is considered as a state. The length of the input vector and the length of the sequence depend on both the size of the window and the distance between two adjacent windows in STFT. These values were obtained in the hyperparameter tuning phase and are listed in the hyperparameters of the model.

All used hyperparameters of both RNN models are summarised in Table 1.

### 2.6 Data processing and performance metrics

As previously mentioned, the extracted dataset consisted of 16,520 samples, of which 10,367 belong to Risso’s dolphin and 6,153 belong to the broadband species. Each sample consists of the click’s waveform and its label. All waveforms were normalised and zero-padded to a length of 512 points.

The dataset was divided into six folds, one of which was separated for hyperparameter tuning. The other five folds were used in the form of a five-fold-cross-validation structure to train and evaluate the models. The models were trained five times independently over about 11,000 samples and evaluated on about 2,750 distinct samples.

To prevent overfitting, the early stopping condition was applied to the training loops using 15% of the training data as a validation set. Training was stopped if the validation loss increased continuously for 30 epochs and according to the validation loss, the best model was returned.

In the training phase, we balanced the dataset by oversampling the broadband species. We used the binary classification loss function and Adam optimisation algorithm. The training dataset was divided into batches containing 16 samples. The Risso’s dolphin class was considered the positive class, and we calculated the following performance metrics:

– **Accuracy**, is a metric that generally describes how the model performed across all classes. It was calculated as the ratio between the number of correct Risso’s dolphin predictions to the total number of click predictions.
– **Precision**, is a metric that measures the model’s accuracy in classifying a click as Risso’s. It was calculated as the ratio between the number of Risso’s clicks correctly classified to the total number of clicks classified as Risso’s (either correctly or incorrectly).
– **Recall**, is a metric that measures the model’s ability to detect Risso’s dolphin clicks. It is calculated as the ratio between the number of correctly classified Risso’s clicks to the total number of Risso’s clicks.
– **F1-score** is the harmonic mean between precision and recall.
– **Binary Cross-Entropy Loss (or Log Loss)** is another metric to assess the performance of a classification problem, and which measures the difference between the two probability distributions.

## 3. RESULTS

Table 2 shows the average performance metrics (accuracy, precision, recall, F1 score, loss) for the four models tested. The individual training and validation loss curves are presented for each model and each fold in Figure S2 (supplementary material). These learning curves demonstrate the performance of the neural network over the iteration while training. The CNN all converged quickly, showing signs of overfitting (Fig. S2A & B). The inflection points on the graph (‘best epoch’) show the point at which training could be halted to avoid overfitting. Similarly, the RNN also showed signs of overfitting, although it took more epochs to achieve training in the case in which raw data was used as input (Fig. S2C & D).

**Table 2:** Performance metrics (accuracy, precision, recall, F1 score) statistics (mean, minimum, maximum and standard deviation) and number of parameters for each model

| metric |  | 1) CNN-raw | 2) CNN-Spectrogram | 3) RNN-raw | 4) RNN-Spectrogram |
| --- | --- | --- | --- | --- | --- |
| **Accuracy** | **max** | 0.9241 | 0.9669 | 0.9571 | 0.8747 |
|  | **mean** | 0.9091 | 0.9567 | 0.9473 | 0.8466 |
|  | **min** | 0.8965 | 0.9452 | 0.9324 | 0.8083 |
|  | **std** | 0.010043 | 0.007827 | 0.010317 | 0.028896 |
| **Precision** | **max** | 0.9468 | 0.975 | 0.9626 | 0.9027 |
|  | **mean** | 0.9347 | 0.9707 | 0.9548 | 0.8735 |
|  | **min** | 0.9054 | 0.9617 | 0.9452 | 0.8297 |
|  | **std** | 0.016755 | 0.005617 | 0.006885 | 0.030346 |
| **Recall** | **max** | 0.9473 | 0.974 | 0.9803 | 0.9074 |
|  | **mean** | 0.9199 | 0.9599 | 0.9615 | 0.8842 |
|  | **min** | 0.8889 | 0.9369 | 0.9473 | 0.8472 |
|  | **std** | 0.023345 | 0.014887 | 0.013296 | 0.024051 |
| **F1 Score** | **max** | 0.9394 | 0.9736 | 0.9663 | 0.8998 |
|  | **mean** | 0.9269 | 0.9652 | 0.9581 | 0.8786 |
|  | **min** | 0.9151 | 0.9554 | 0.9462 | 0.8512 |
|  | **std** | 0.008621 | 0.006602 | 0.008335 | 0.022133 |
| **Loss** | **max** | 0.5247 | 0.4849 | 0.4954 | 0.5692 |
|  | **mean** | 0.5159 | 0.4790 | 0.4852 | 0.5442 |
|  | **min** | 0.5090 | 0.4730 | 0.4802 | 0.5257 |
|  | **std** | 0.005615 | 0.004534 | 0.006533 | 0.018491 |
| **Number of parameters** |  | 4,126,833 | 10,835,985 | 18,541 | 47,561 |

The average performance metrics and the number of parameters used for each model are presented in Table 2. The individual performance metrics for each model and each fold are illustrated in Figure S3. All models showed high performance, with almost all metrics above 90 %. The CNN fed with the spectrograms and the RNN fed with raw data both showed a precision, accuracy, recall and F1 score of more than 95 %. This indicates that these models achieved many correct positive classifications while minimising false positives. A good balance between precision and recall is indicated by high F1 scores (97 % for both models). We used binary cross-entropy (log loss) as our loss function. As the classes were unbalanced (prevalence 63%) the log loss metrics for both models also showed good performance (0.48 and 0.49, respectively).

## 4. DISCUSSION

We demonstrate here the effective application of deep neural networks for classifying delphinid clicks between Risso’s dolphins and two broadband species (common or bottlenose dolphins). CNN and RNN models were trained on manually labelled clicks and generalised well to all clicks recorded in the dataset derived from an environment with high tidal current speeds and an MRE infrastructure in place and thus varying background noise. Both CNN and RNN automatically learned the representations needed from the input raw data and the spectrograms, bypassing the pre-processing phase. Feature extractions were thus not required in the case where the raw data was fed to the models, and clicks could be classified based on their raw waveforms by the CNN and the RNN with accuracy, precision and recall higher than 90 %. The highest performance was achieved by CNN fed with spectrograms (F1 score 97 %) and the RNN fed with raw data (F1 score 96 %).

While CNN with spectrograms give a slightly better performance result, we prefer and recommend the RNN model with raw data, taking into account the lower number of parameters that is preferable in terms of operability and processing effort. The performance of the RNN confirms the ability of recurrent models to process and understand consecutive data. Furthermore, the choice of RNN with raw data avoids any data pre-processing procedures, making the workflow simpler than when using spectrograms.

While artificial neural networks have already been successfully used to classify odontocete clicks between sperm whales and long-finned pilot whales (Jiang et al., 2018), this is the first application of supervised neural networks to classify delphinid clicks into taxonomic groups. Delphinid clicks are particularly challenging to classify due to their high variability, notably due to variable source levels and beam width, and rapid attenuation of the clicks (Au and Benoit-Bird, 2003; Finneran et al., 2016; Kloepper et al., 2012; Moore et al., 2008). Only nonsupervised networks have so far been tested successfully to classify delphinid clicks (Frasier, 2021; Frasier et al., 2017). Although this methodology was successful in separating group clicks into different classes, only one taxonomic group of delphinid was clearly identifiable (Risso’s dolphin) in the latest study (Frasier, 2021). Here, we show that supervised networks allow click classification from taxonomically different species.

Traditionally, acoustic recordings are taken in conditions where the background noise is limited or can be filtered out. However, noise is sometimes inevitable and may exhibit elevated variability, for example in a region prone to storms, near a submerged anthropogenic structure producing noise, or as in this study, in strongly tidal sites where current speed fluctuates considerably and includes an inherently noisy MRE infrastructure. Our results show that neural networks are resilient to varying levels of background noise and offer good generalisation performance. Therefore, such an automatic classifier could be used in coastal regions with strong tidal currents as well as sites with MRE infrastructures presenting varying levels of background noise (see e.g., Benjamins et al., 2017; Gillespie et al., 2021; Palmer et al., 2021).

This study focused on the classification of two groups of delphinid species in one region, using one month of PAM data. The classifier could be generalised by including clicks detected by the automatic detector produced by additional background noise sources, such as snapping shrimps or other organisms and anthropogenic clicks derived from maritime transport sources or noise derived from other types of undersea infrastructure. Also, clicks from a certain species can sometimes present different features and can be subclassified interspecifically, as done for the sperm whale *Physeter macrocephalus* with artificial networks (Schaar et al., 2009, 2007). Ecological insights may thus be gained from being able to distinguish click patterns within a species with non-supervised neural networks fed with raw data. Furthermore, as clicks are usually emitted in groups, or ‘click trains’ (Au, 1993, 2002), they could be fed in this form to the machine, rather than individually, so that the temporal and spectral features of the click trains may be considered, which may improve the machine learning model performance. More generally, future research should explore the use of such models to classify echolocation clicks from other species, recorded in different geographic regions and over different seasons. Deep neural networks have previously been shown to generalise well to recordings taken at different spatial and temporal scales, even if not represented in the training data (Shiu et al., 2020).

Advances in automatic sound classification allow rapid processing of large volumes of data. The classification of marine mammal clicks is especially valuable for research to improve understanding and management of marine ecosystems. With the increase in anthropogenic noise in the ocean (Duarte et al., 2021) and the development of underwater energy sources, it is especially important to monitor the presence of these animals despite the presence of background noise. Furthermore, although PAM data is commonly analysed retrospectively, there are emerging policy requirements to develop real-time *in situ* monitoring and classification of marine mammals to estimate collision risk as new tidal stream MRE infrastructures are developed and operated. Our work demonstrates that state-of-the-art neural networks can successfully achieve classification tasks under these conditions.

## DATA AVAILABILITY STATEMENT

The trained machine learning model is openly available at XXXXXXXXX. Further inquiries can be directed to the corresponding author.

## ACKNOWLEDGMENTS

This work was funded by the SEACAMS2 and SEEC (Smart Efficient Energy Centre) projects, both part-funded by the European Regional Development Fund through the Welsh Government.

